# Elemental profiling and genome-wide association mapping reveal genomic variants modulating ionomic composition in *Populus trichocarpa* leaves

**DOI:** 10.1101/2024.05.03.592412

**Authors:** Raphael Ployet, Kai Feng, Jin Zhang, Ivan Baxter, David C. Glasgow, Jin-Gui Chen, Gerald A. Tuskan, Timothy J. Tschaplinski, David J. Weston, Madhavi Z. Martin, Wellington Muchero

## Abstract

The ionome represents elemental composition in plant tissues and can be an indicator of nutrient status as well as overall plant performance. Thus, identifying genetic determinants governing elemental uptake and storage is an important goal in plant breeding and engineering. In this study, we coupled high-throughput ionome characterization with high-resolution genome-wide association studies (GWAS) to uncover genetic loci that modulate ionomic composition in leaves of 584 black cottonwood poplar (*Populus trichocarpa*) genotypes. Congruence of alternate ionomic profiling platforms, i.e., inductively coupled plasma-mass spectrometry (ICP-MS), neutron activation analysis (NAA) and laser-induced breakdown spectroscopy (LIBS), was performed on leaf samples from a subset of the population. Significant agreement was observed across the three platforms with some notable exceptions for individual elements. Subsequently, we used the ICP-MS platform to profile the 584 genotypes focusing on 20 elements. GWAS performed using a set of high-density (>8.2 million) single nucleotide polymorphisms (SNP), identified multiple loci significantly associated with variations in these mineral elements. The potential causal genes for variations in the ionome were significantly enriched in genes whose homologs were previously associated to ion homeostasis in other species. Notably, a polymorphic copy of the high-affinity molybdenum transporter MOT1 was found directly associated to molybdenum content in leaf tissues. The results of the GWAS also provided evidence of physiological and genetic interactions between mineral elements in poplar. The new candidate genes predicted to play a key role in cross-homeostasis of multiple elements are new targets for engineering a variety of traits of interest in tree species.

## 1. Introduction

A multi-omics profile includes representation by four biochemical pillars, i.e., the transcriptome, proteome, metabolome, and ionome, which collectively describe, in part, the molecular functioning of a living organism. The study of the transcriptome (Klepikova et al., 2016; Zhu and Wang, 2000), proteome (Cánovas et al., 2004; Abraham et al., 2015; Mergner and Kuster, 2022; Castillejo et al., 2023), and metabolome (Fiehn et al., 2000; Tschaplinsky et al., 2019) have been undertaken in substantial detail in many organisms, but the ionome investigations are just emerging (Rea, 2003; Baxter, 2009; Singh et al., 2013) and many of the genes and gene networks involved in ionome regulation remain largely unknown. The ionome includes all metal, metalloid, and nonmetal elements present in an organism (Salt et al., 2008), including all macro- and micro-nutrients. In plants, macro-nutrients (N, P, K, Ca, S, Mg) are required in large amounts, whereas micro-nutrients and trace elements (Ni, Mo, Cu, Zn, Mn, B, Fe, Cl) are usually required in smaller amounts (Williams and Salt, 2009). Macro-nutrients play many pivotal roles in plant development. For example, K and Ca are the most abundant free cations in plant cells (Sardans and Peñuelas, 2021), where K regulates activity of numerous enzymes, playing a role in photosynthesis and biosynthetic pathways and is involved in osmo-regulation (Cui and Tcherkez, 2021). It allows plant cells to regulate their water potential which in turns allows plants to control their stomatal aperture, and uptake water and nutrients, contributing to development and adaptation to water deficit conditions (Cakmak, 2005; Mostofa et al., 2022). For most plant species, abundance of K was shown to greatly improve primary and secondary growth (Sardans and Peñuelas, 2021). Ca is also involved in a range of developmental processes, including plant cell walls formation through its role in pectins organization and as an intra-cellular messenger that allows plants to respond to environmental stimuli. Another essential macronutrient, P, is incorporated in numerous biomolecules, such as nucleic acids and phospholipids playing structural roles and in signaling via protein phosphorylation. In addition to these major elements, micronutrients play key roles as enzymatic co-factors and signaling molecules. While macro-nutrients are generally growth limiting factors in case of their deficiency, a severe impact of depletion of other micronutrients, such as Zn and Mo, has also been documented, leading to impaired growth and poor resistance to stresses (Arnon and Stout, 1939; Fageria et al., 2002). Mo is an example of micronutrient required in trace amounts, serving as a co-factor for numerous molybdoenzymes that catalyze oxidation–reduction reactions essential for nitrogen and sulfur metabolism (Tejada-Jimenez et al., 2013).

Other elements such as Na, Co, Al, and Se are known to have biological functions in plants (Williams and Salt, 2009). For example, Na, a non-essential element for most plants, is also one of the most abundant ions in plant tissues (Subbarao et al., 2010). While it is mostly known for its toxic effect and negative impact on growth at high concentration (Adams and Shin, 2014; Maathuis, 2014), it was shown that amendment with Na could at least partially alleviate K depletion through its unspecific function as a monovalent cation, having a positive effect on growth and resistance to water deficit in different species (Epron et al., 2012; Erel et al., 2014; Favreau et al., 2019; Tomemori et al., 2002; Wakeel et al., 2011, 2010). This highlights that different ions can have overlapping functions, sometimes competing for transport or simply showing strong interactions at chemical, physiological and/or genetic levels (Baxter, 2015; Bouain et al., 2019).

In addition to the uptake, compartmentalization of these elements is a key regulatory process. Ion transporters are of central importance, making these proteins the primary focus of most research involved in characterizing the underlying genetic components of ionome regulation in plants. However, ion homeostasis is often regulated by a complex cooperation of multiple genes and the existence of different levels of interactions between mineral elements make ionome investigation challenging for conventional forward genetics approaches (Mäser et al., 2001).

Watanabe et al., (2007) used neutron activation analysis (NAA) to analyze 44 elements in 2,228 leaf samples from 670 species of terrestrial plants spread across 138 distinct families. They showed that over 25% of the total variation in leaf elemental composition could be assigned to the phylogenetic family level, (Watanabe et al., 2007), suggesting inheritance and a genetic component to variation in the ionome. Similar observations were made by Neugebauer and colleagues when surveying many studies performed in angiosperms (Neugebauer et al., 2018). The fact that these phylogenetic effects explained not only variations in the macronutrients N and P, but also other micronutrients and trace elements, support the hypothesis that a large proportion of the ionome in plants is under tight genetic control. While several studies have successfully captured variations of the ionome across populations of dicots (*Arabidopsis*, potato, and Cassava; Lahner et al., 2003; Baxter et al., 2012; Yin et al., 2022) and monocots (rice and maize; Yang et al., 2018; Wu et al., 2021), very little is known about variations in the mineral content of populations of perennial woody species (Singh et al., 2013).

The use of *Populus* spp. (poplars, aspens, and cottonwoods) is increasing globally as demand for lignocellulosic biomass increases (Clifton-Brown et al., 2019). The *Populus* genus, consisting of woody perennial species that occupy diverse habitats ranging from the tropics to the sub-arctic, are dominant members of riparian systems and are important pioneer species within boreal forests. *Populus* is largely undomesticated with natural populations exhibiting a high degree of genetic and phenotypic trait variability. Variation in mineral accumulation in natural populations of poplars and the genetic determinants of variation remain mostly unknown. As an alternative to conventional genetic screens, genome-wide associations studies (GWAS) have been instrumental for discovering genetic determinants of complex traits in many plant species, including poplar (Tuskan et al., 2018a; Gupta et al., 2019). By precisely linking genomic mutations to population-wide phenotypic variation, GWAS performed in poplar pointed to regulatory loci for a variety of phenotypes, including phenology (Evans et al., 2014), lignin biosynthesis (Xie et al., 2018), metabolite biosynthesis (Zhang et al., 2018) and disease resistance (Muchero et al., 2018), leading to the discovery of new targets for genetic engineering. GWAS approaches have also been successfully applied to the discovery of regulators of the ionome in annual plants (Lahner et al., 2003; Baxter et al., 2010; Forsberg et al., 2015; M. Yang et al., 2018; Wu et al., 2021; Yin et al., 2022).

In most cases, the main challenge for performing GWAS is to have a cost-effective, sensitive, and scalable method to analyze the phenotype of interest. For ionomics, the three most common methods for high-throughput elemental analysis are: 1) atomic absorption spectroscopy (AAS), 2) inductively coupled plasma-optical emission spectroscopy (ICP-OES), and 3) ICP-mass spectrometry (ICP-MS). In addition to these, LIBS has also been used as a novel high-throughput technique to determine the ionome of poplar leaves (Kaiser et al., 2012; Krajcarova et al., 2013; Martin et al., 2017), while X-ray fluorescence spectroscopy (XRF) has been used for elemental analysis of multiple plant samples (Salt et al., 2008).

In this work, we systematically assessed the sensitivity, and scalability of multiple platforms for characterizing the elemental profile of a large number of poplar leaf samples. Using ICP-MS, we then profiled the ionome of a large population of natural variants of black cottonwood poplar (*Populus trichocarpa*) grown in a common garden. This allowed investigating to what degree genetic variation impacts poplar leaf elemental composition. Then, the profiles of 19 individual mineral elements were used as quantitative traits in a GWAS, to identify the genetic determinants underlying leaf elemental composition in poplar. This approach identified several known key regulators of elemental accumulation in plant tissues and several novel candidate genes that play potential critical roles in ion homeostasis and in the interaction between mineral elements, providing new targets for engineering traits such as improved growth and resistance to biotic and abiotic stressors.

## 2. Material and Methods

### 2.1. Plant material and sample collection

A population of 1,089 black cottonwood genotypes (*P. trichocarpa*) was assembled from native stands to encompass the central portion of the natural range of the species, stretching from 38.8° to 54.3° N latitude from California, USA to British Columbia, Canada. Establishment of the common garden, growth conditions, and site maintenance have been described by Muchero et al. (Muchero et al., 2015). In this study, leaf samples for ionomic profiling were collected from 4-year-old trees, from a field located in Clatskanie, Oregon, USA (46°6′11″N 123°12′13″W). A subset of 584 out of the 1,089 *P. trichocarpa* genotypes were represented in this sampling. A single fully mature (LPI 7-9) leaf on the south side of the tree exposed to full sunlight conditions was removed from the tree within a 6-hour window centering on solar noon and immediately frozen under dry ice before processing.

### 2.2. Sample preparation

Initially, 25 poplar samples and 5 NIST standard reference materials were chosen for LIBS and NAA. The 25 poplar leaf samples were finely ground to 40 μm particle size. To obtain consistent pellets that would be robust enough to withstand the laser energy that is used to excite the samples, 75 mg of the ground powder for each sample was weighed then humidified by equilibrating the weighed samples in a chamber with open beakers of water for 12 hours. The humidified powder was pelletized for 3 min at pressures of 2000 psi in a Carver press with a 6 mm evacuable die (Martin et al., 2010). Five NIST standard reference materials, including spinach leaves, orchard leaves, tomato leaves, pine needle, and bovine liver, were used as comparators. Two sets of 30 calibration samples were prepared. One set was used to perform the LIBS measurements and the second set was used to do the NAA experiments. No chemical treatment was required for either technique. The detection of the 9 elements (Ca43, Mg25, Mn55, Na23, K39, Cu, Fe58, Cl35, and Al27) using LIBS technique were correlated to the NAA results and used to construct a regression model for these elements.

### 2.3. Laser Induced Breakdown Spectroscopy (LIBS)

Previously, LIBS spectra were correlated to NAA concentrations for Ca, Mg, Mn, Al, Cu, and K (Martin et al., 2015, 2017). In this study, additional elements have been identified using LIBS and correlated to the standard NAA technique. These elements were Na23, Fe57, Cr52, Co, As75, Zn66, Br80, Rb85, Sb119, La139, Ce140, and Sm150. The excitation laser was a Q-switched Nd:YAG laser with frequency doubled output wavelength of 532 nm and a laser energy of 45 mJ (Martin et al., 2010). A displacement laser (650 nm) was employed to maneuver the vertical position of the upper surface of the sample, ensuring that the excitation laser beam remained focused and that the plasma was optimally located with respect to the collection lenses. The light emitted by the plasma at the focal volume was collected by a set of fused silica lenses and re-focused into a low O-H silica fiber of internal diameter 100 μm. This optical fiber delivered the light from the plasma spark to an Echelle spectrometer (Catalina Scientific model SE 200). A high-order dispersion module was used to separate the white light formed in the plasma into respective wavelengths in the large wavelength range (190-800 nm) with a wavelength resolution of 0.06 nm in the optimized range of 300-500 nm. An intensified charge coupled detector (ICCD) (Andor Technologies, Belfast, Northern Ireland) was used to convert the optical signal into a digital signal. The data for all pellets were collected using a 1 μs delay and 10 μs emission detection window. The laser was operated at 10 Hz repetition rate, which allowed data collection every 100 ms.

### 2.4. Neutron Activation Analysis (NAA)

The NAA of leaf samples was described in a previous study (Martin et al., 2017), and a similar protocol was used for this study, with minor modifications. The irradiations were made in the pneumatic tube PT-1, for a duration of 300 s, and counting followed 5-7 days later. Measurement of the samples on high-purity germanium (HPGe) detectors occurred both at 30 mm shelf height and sometimes directly on the endcap. Because comparator NAA was employed throughout this work, the effects of efficiency calibration, coincidence summing, gamma-ray attenuation, and the uncertainty of nuclear parameters, such as cross sections and gamma-ray branching ratios, were assumed to be constant.

### 2.5. Inductively Coupled Plasma Mass Spectrometry

Leaf samples from the 584 *P. trichocarpa* genotypes were analyzed for ionomic composition using ICP-MS. In total, 20 elements were profiled, including Aluminum (Al27), Arsenic (As75), Boron (B11), Cadmium (Cd111), Calcium (Ca43), Cobalt (Co), Copper (Cu), Iron (Fe57), Magnesium (Mg25), Manganese (Mn55), Molybdenum (Mo), Nickel (Ni60), Phosphorus (P31), Potassium (K39), Rubidium (Rb85), Selenium (Se82), Sodium (Na23), Strontium (Sr88), Sulfur (S34), and Zinc (Zn66), following a protocol adapted from Baxter et al. (2008a). For subsequent analyses, the quantifications were converted to total element concentration, and isotope numbers are not represented.

### 2.6. GWAS and candidate gene identification

The GWAS were performed with elemental composition as the phenotypic variable by using the analytical platform described by Zhang et al. (Zhang et al., 2018). Briefly, the single nucleotide polymorphism (SNP) and indel dataset used in this study is available at http://bioenergycenter.org/besc/gwas/. To assess genetic control, we used the EMMA algorithm in the EMMAX software with kinship as the correction factor for genetic background effects to compute genotype to phenotype associations based on 8,253,066 SNP variants with minor allele frequencies ≥0.05 identified from whole-genome resequencing. For all phenotypes, the top 100 most significant SNPs (lowest *p*-value) with -log10(*p*) >6 were retained for further analysis. For each significant SNP, the flanking genes detected in a 20-kb window centered around the SNP position, were considered as potential candidate genes for that locus. Results were visualized as networks using Cytoscape v3.9.1.

Co-expression data was obtained by calculating pairwise Pearson correlations between gene profiles across 8 tissues of *P. trichocarpa* (total of 23 RNAseq samples) from previously published data (SRA accession PRJNA320431; Shi et al., 2017). RNAseq data were retrieved from SRA and mapped to *P. trichocarpa* genome v3.0 using a similar approach as described by Sundell et al. (Sundell et al., 2017). Briefly, quality of the reads was assessed using FastQC (http://www.bioinformatics.babraham.ac.uk/projects/fastqc/ v0.11.9), then residual adapters and low quality reads were trimmed using Trimmomatic v0.39 (Bolger et al., 2014), and filtered reads were mapped to the reference genome using STAR v2.7.6 (with parameters – outFilterMultimapNmax 100; Dobin et al., 2013). Transcript per million (TPM) values were calculated using Stringtie v1.3.4 (with parameter -M 0.75; Pertea et al., 2015). Significant Pearson correlations with False Discovery Rate (FDR) adjusted *p*<1e-3 were retained for each pair of genes. For each potential candidate gene identified by the GWAS, the top 500 most highly co-expressed genes (highest absolute r^2^ and FDR <1e-3) were selected, and GO enrichment was performed considering the GO annotation of the first BLAST hit in *Arabidopsis* of the *P. trichocarpa* gene (*P. trichocarpa* genome annotation file v3.0, available at Phytozome V12), using the R package TopGO. GO terms with FDR adjusted *p*≤0.05 were considered significantly enriched and reported in the results.

Protein alignment of AT2G25680 (AtMOT1), Potri.006G245900 (PtrMOT1-2) and Potri.006G246000 (PtrMOT1-1) were generated using the online multiple sequence alignment tool Clustal Omega (https://www.ebi.ac.uk/Tools/msa/clustalo/; Madeira et al., 2022) with default parameters, and visualized using Jalview (Waterhouse et al., 2009).

## 3. Results

### 3.1. Comparisons of LIBS, NAA, and ICP-MS platforms for elemental profiling of poplar samples

In order to identify the most suitable platform (i.e., the ability to detect a wide range of elements, high throughput, and accurate) for the profiling of mineral elements in poplar leaf samples of a large natural population, a subset of 25 samples (out of 584 samples) was analyzed using three different methods and results were compared.

Rapid multi-elemental microanalysis of different types of sample matrices (solid, liquid, gases, and aerosols) was previously demonstrated using the LIBS technique (Cremers and Radziemski, 2017; Radziemski and Cremers, 2020). The detection of most elements from these matrices was in the parts-per-million (ppm) range, with the resulting spectrum consisting of emission peaks that are representative of all the elements present in the sample. Under our conditions, LIBS spectra were able to differentiate elemental peaks among the four NIST standards (Fig. 1a). Based on these results a variety of major elements were detected in the spectra of the 25-member subset of poplar leaves. Elemental concentrations varied significantly among the tested genotypes, as exemplified in Fig. 1b with profiles of three genotypes that showed low, medium, and high levels of certain elements.

**Figure 1.**
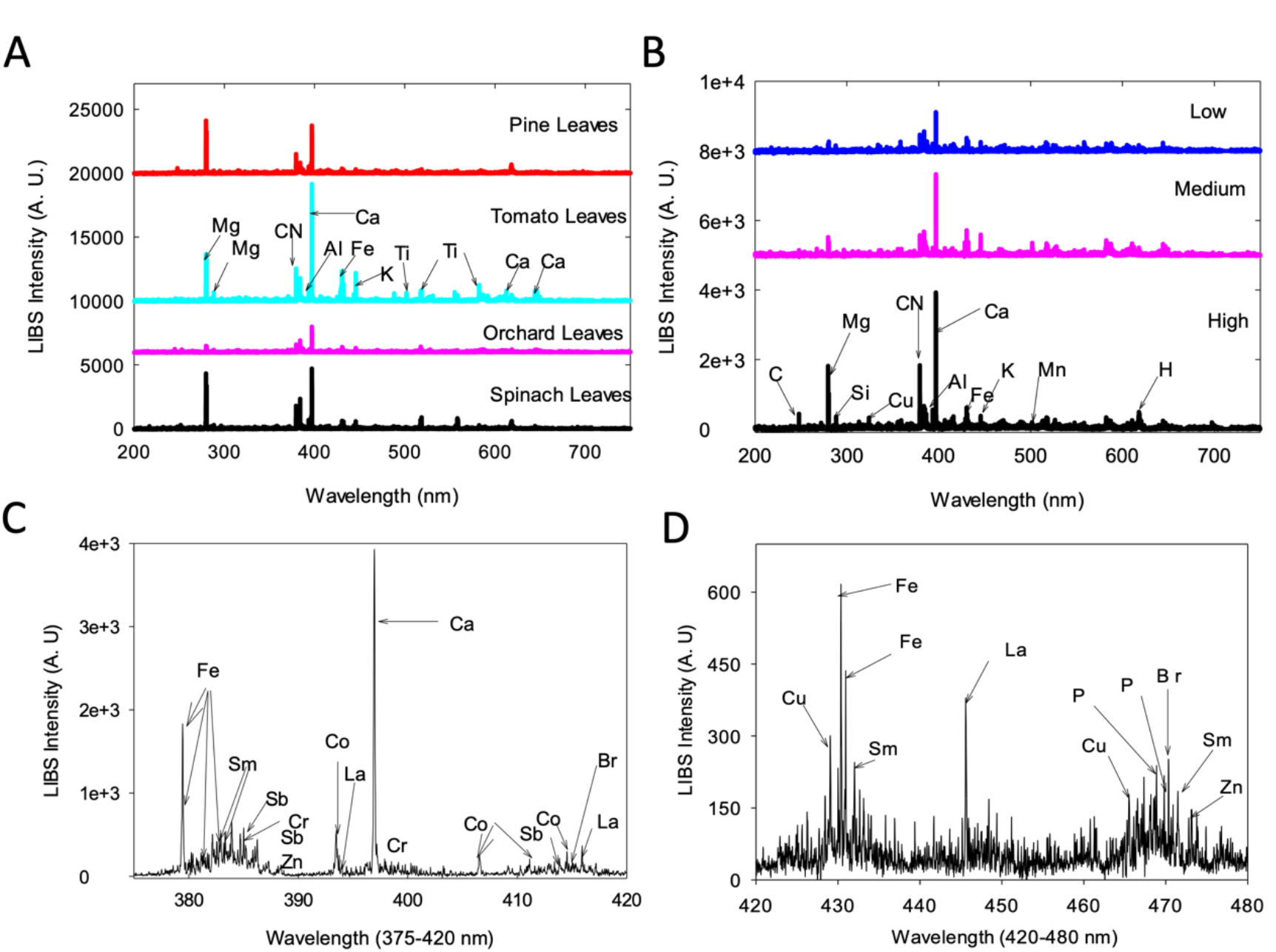
LIBS spectra obtained for NIST standards and poplar leaf samples. **A**) LIBS spectra for four NIST standards, **B**) LIBS spectra for 3 poplar samples with low, medium, and high (top to bottom) concentrations of elements, **C**) LIBS spectrum for a poplar leaf sample (high concentration) that shows 9 minor elements present in the sample in a narrow spectral window of 375-420 nm and **D**) the detection of copper in the window 420-480 nm.

NAA is a nuclear analytical trace element metrology technique that utilizes neutron irradiation to induce radioactivity that is then emitted and detected to reveal the number and kind of trace elements composing a sample. Results of quantification of Ca, Mg, Mn, Al, Cu, and K from poplar leaves through NAA were highly correlated to what were obtained through LIBS (Martin et al., 2017). However, the LIBS method was able to detect a suite of minor elements that were not available with the NAA method, including Sm, Sb, Cr, Co, La, and Br (Fig. 1c), with Cu additionally detected in the window 420-480 nm (Fig. 1d).

ICP-MS is a versatile trace element quantification method that is well-suited to large sample sets because it can detect many of the elements simultaneously. In contrast to LIBS and NAA, sample dissolution is required for ICP-MS, which is often carried out in acid using microwave-assisted digestion. We compared the NAA results to both LIBS and ICP-MS for trace and matrix elements that were common to the three techniques (Table 1, Fig. 2). The profiles of several elements showed high correlation between all three methods (e.g., Mg, Ca, Mn; r^2^>0.7, Fig. 2a), indicating that these elements are relatively homogeneous in the samples and that the three tested platforms perform equally well (Table 1). Al and Zn quantification using ICP-MS and NAA showed lower correlation (r^2^ = 0.49; Fig. 2b, Table 1), while quantifications for Cu, Na, and Ru were poorly correlated for these two methods (r^2^ ranging from 0.001 to 0.130; Fig. 2c, Table 1). The results obtained for K were also poorly correlated across methods (Fig. 2d), suggesting that sample preparation and the method itself affect the quantification of mineral elements.

**Figure 2.**
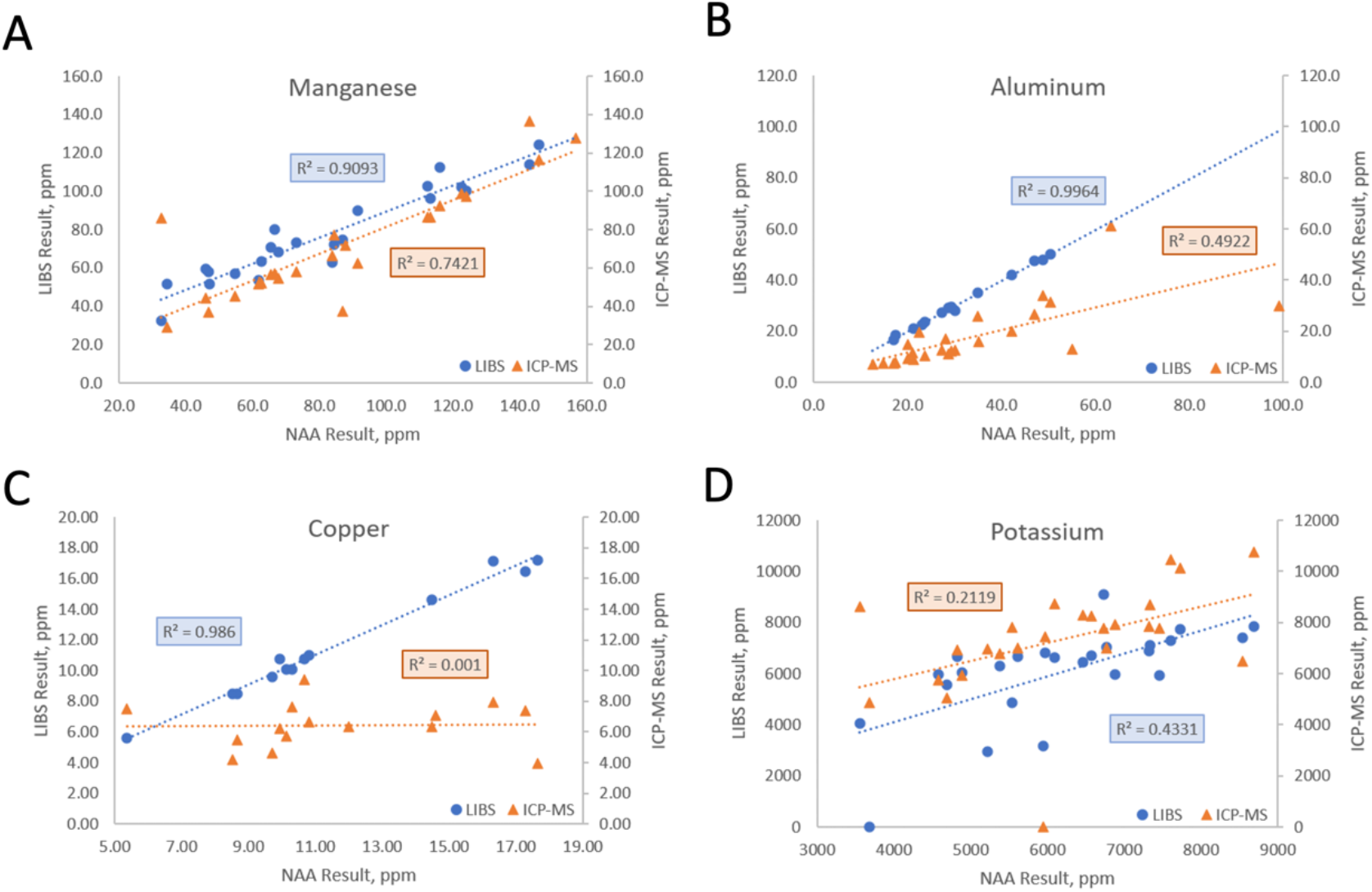
ICP-MS and LIBS comparison to NAA for Mn, Al, Cu, and K. The linear correlation coefficients reflect the level of agreement between values obtained from different analytical platforms for these four elements quantified in 25 poplar leaf samples.

**Table 1.** NAA, ICP-MS, and LIBS Pearson Correlation Coefficients. The reactions used for NAA quantification of the elements are also presented.

| Element | NAA / ICP-MS<br>Correlation, $r^2$ | NAA / LIBS<br>Correlation, $r^2$ | NAA<br>Reaction |
| --- | --- | --- | --- |
| Mg | 0.762 | 0.776 | $^{26}\text{Mg} (n,\gamma) ^{27}\text{Mg}$ |
| Ca | 0.882 | 0.894 | $^{48}\text{Ca} (n,\gamma) ^{49}\text{Ca}$ |
| K | 0.395 | 0.351 | $^{41}\text{K} (n,\gamma) ^{42}\text{K}$ |
| Na | 0.034 | 0.764 | $^{23}\text{Na} (n,\gamma) ^{24}\text{Na}$ |
| Mn | 0.742 | 0.909 | $^{55}\text{Mn} (n,\gamma) ^{56}\text{Mn}$ |
| Cu | 0.001 | 0.986 | $^{65}\text{Cu} (n,\gamma) ^{66}\text{Cu}$ |
| Al | 0.492 | 0.996 | $^{27}\text{Al} (n,\gamma) ^{28}\text{Al}$ |
| Fe | 0.687 | 0.984 | $^{58}\text{Fe} (n,\gamma) ^{59}\text{Fe}$ |
| Zn | 0.491 | 0.990 | $^{64}\text{Zn} (n,\gamma) ^{65}\text{Zn}$ |
| Co | 0.831 | | $^{59}\text{Co} (n,\gamma) ^{60}\text{Co}$ |
| As | 0.778 | | $^{75}\text{As} (n,\gamma) ^{76}\text{As}$ |
| Rb | 0.130 | | $^{85}\text{Rb} (n,\gamma) ^{86}\text{Rb}$ |

### 3.2. Elemental profiling of a population of poplar natural variants and GWAS

Since the IPC-MS method had a broad range of detectable elements, a relatively high throughput, and showed significant correlation with other methods for most elements, we used the IPC-MS method to characterize the profile of 20 elements (Fig. 3) across the 584 genotypes of the population. We observed significant variation in the content of all the elements. Interestingly, macronutrients essential for plant growth (P, K, Ca, and Mg) had relatively smaller standard deviations, ranging from 12 to 27% within the population, than those of most micronutrients (Fe, Zn, Cu, Mn, B, Ni, and Mo), which varied between 32 and 50% (Supplementary Data S1). Other trace elements, such as Cd, As, and Co, also showed larger variation than macronutrients (relative standard deviation ranging from 41 to 45%), which is consistent with observations from previous studies in other species (M. Yang et al., 2018).

**Figure 3.**
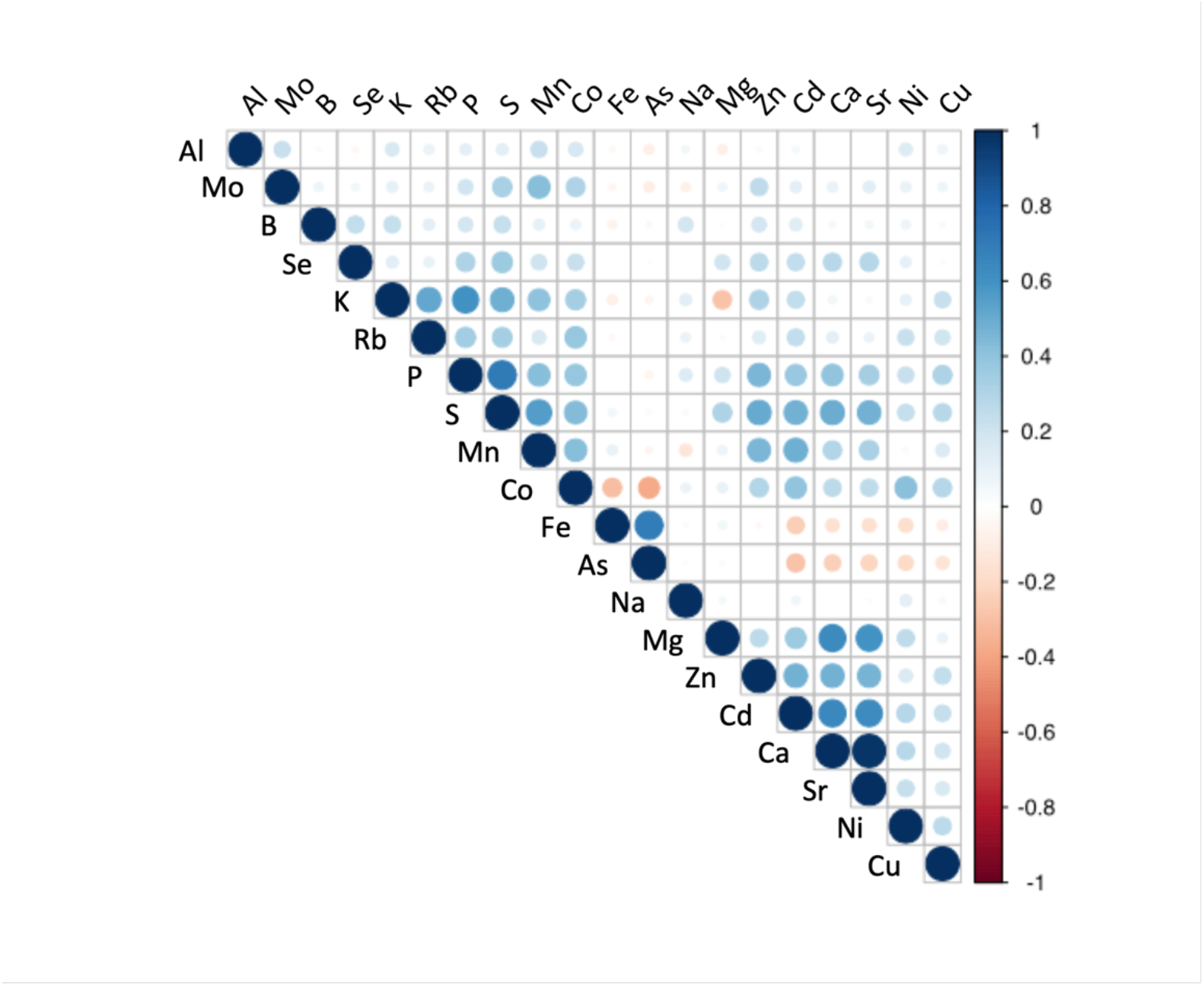
Correlation analysis for leaf elemental composition across 584 *Populus trichocarpa* genotypes. Pearson pairwise correlation coefficients were calculated from the profile of each element across the 584 individual genotypes.

Eight elements exhibited high positive correlation in their abundances, suggesting a possible shared genetic regulation and/or interactions in common biological processes. These elemental pairs had significant Pearson correlation values greater than 0.60 (*p* ≤0.05; Fig. 3) and included: Ca/Sr (r^2^=0.97), P/S (r^2^=0.72), As/Fe (r^2^=0.69), Ca/Cd (r^2^=0.65), Ca/Mg (r^2^=0.64), Cd/Sr (r^2^=0.63), Mg/Sr (r^2^ =0.60) and K/P (r^2^=0.60).

The GWAS was performed to test associations between >8.2 million SNPs and profiles of 19 elements across 584 individual genotypes. Since boron had a distribution that significantly deviated from a normal distribution, it was not considered in the GWAS (Fig. S1a,b). The GWAS performed on the ICP-MS generated ionomic profiles, identified 631 SNPs significantly associated with at least one of the 19 elements at a threshold of -log10 (*p*) >6 (Supplementary Data 2; Supplementary Fig. 2). Of the 19 elements, Al, Cu, and Na accounted for 69% of all associations detected, including associations with a total of 434 unique SNP positions (207, 114, and 113 SNPs, respectively), suggesting that Al, Cu, and Na content are highly polygenic traits in poplar (Supplementary Data 3). Since the comparison of the analytical methods ICP-MS / NAA revealed the potential for more technical variability in the estimation of these three mineral elements, only the strongest associations were considered for further investigation, by considering the top 100 SNPs with lowest *p*-value. Across all other elements, an average of 13 SNPs were significant; for P, K, Ca, and Mg, 3 to 18 SNPs were found significantly associated with variations in macronutrients content (Supplementary Data 3).

### 3.3. Genetic Determinants of Trace Elemental Composition

The 497 SNPs that passed the statistical threshold were associated with 922 potential candidate genes flanking these positions. Among these, 26 genes were putative homologs of *Arabidopsis* genes annotated within at least one of the biological process GO terms ‘multicellular organismal-level chemical homeostasis’ (GO:0140962), ‘intracellular chemical homeostasis’ (GO:0055082), ‘inorganic ion homeostasis’ (GO:0098771), ‘monoatomic ion homeostasis’ (GO:0050801), ‘sequestering of metal ion’ (GO:0051238), ‘regulation of cellular localization’ (GO:0060341) and ‘regulation of sequestering of calcium ion’ (GO:0051282), which represents a significant enrichment of 1.53 for these categories (*p* =0.0132). Notably, 18 of the 922 genes had a putative *Arabidopsis* homolog annotated within the biological process GO term ‘inorganic ion homeostasis’ (GO:0098771), giving a significant enrichment of 1.61 (*p* =0.0187) (Fig. 4, Supplementary Data 4). When filtering the GWAS results at a more stringent threshold on SNP *p*-value (i.e., -log10(*p*)>8), enrichment in genes annotated in that GO term ‘inorganic ion homeostasis’ increased to a value of 2.29 (*p* =0.0428), strongly suggesting that the GWAS was able to capture relevant candidate genes that control variations in elemental concentrations in poplar leaf tissues.

**Figure 4.**
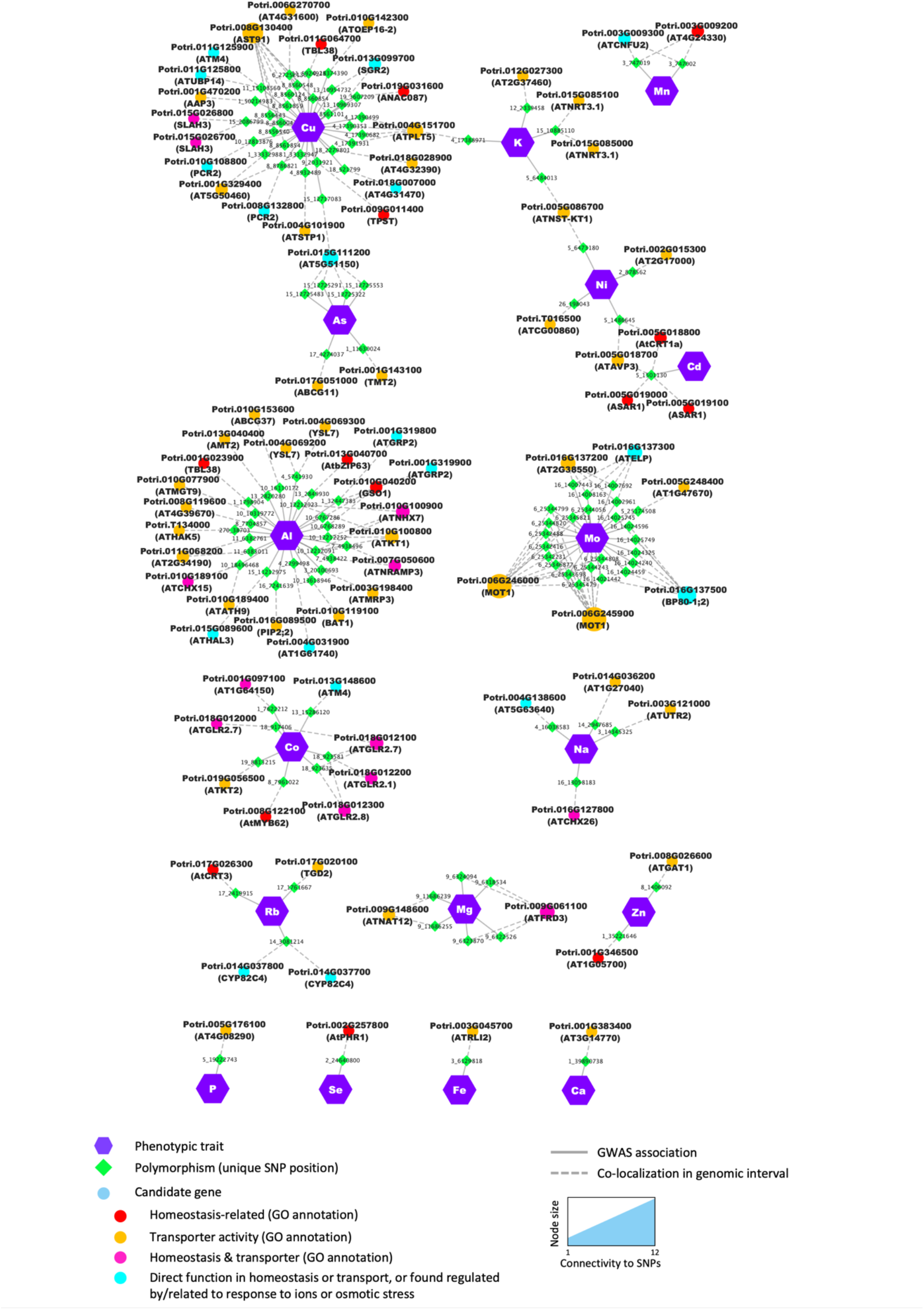
Candidates identified through GWAS were significantly enriched in genes from functional categories related to ion homeostasis and/or with transport molecular function. Nodes (i.e., circles and polygons) represent either phenotypic traits (mineral elements), polymorphisms (unique SNP positions) or potential candidate genes detected in the genomic interval flanking the significant SNP. For each candidate gene, the gene name of the best BLAST hit is provided in brackets in addition to the poplar accession number. When no *Arabidopsis* name was available, the gene accession number in *Arabidopsis* was provided. Edges (i.e., lines) represent the significant association detected by the GWAS (i.e., solid lines) or the colocalization of the SNP and the annotated gene in the genomic interval (i.e., dashed lines). For candidate genes, node color annotates the different functional categories and molecular functions.

Notable candidate genes examples include members of the cation–proton antiporters (CPA) family previously shown to regulate cation and pH homeostasis in plants (Chanroj et al., 2012; Khan et al., 2018). For Na and Al several candidates in the CPA family included Potri.016G127800 and Potri.010G189100, putative homologs of *Arabidopsis* AtCHX26 and AtCHX21/23 (Chanroj et al., 2012), respectively, and Potri.010G100900, homolog of the Na+/H+ exchanger AtNHX7/SOS1 (CPA1) shown to be essential for salt tolerance in *Arabidopsis* (Wu et al., 1996) (Fig. 4, Supplementary Data 4). Potri.015G085000 and Potri.015G085100 homologs of the *Arabidopsis* high-affinity nitrate transporter AtNRT3.1 (Kawachi et al., 2006) were found associated with K content, and Potri.010G108800 and Potri.008G132800, potential homologs of AtPCR2 involved in zinc transport and detoxification in *Arabidopsis* (Song et al., 2010), were found associated with the micronutrient Cu content.

Additionally, 55 genes had their *Arabidopsis* homologs annotated with the molecular function GO term ‘transporter activity’ (GO:0005215; Fig. 4). Although the proportion of genes discovered from that category over the number of genes expected by chance was favorable (enrichment of 1.21), it was not statistically significant (*p* =0.0634), suggesting that although several transporters appear to control elemental composition, as expected, other classes of proteins must be involved in ion homeostasis in poplar leaf tissues. In addition, there were a number of genes predicted to be involved in the direct control of ion fluxes across membranes, including several transcription factors such as Potri.008G122100, homolog of AtMYB62, involved in the response to P starvation in *Arabidopsis* (Devaiah et al., 2009).

Among the candidate genes with transporter activity, one striking example is the detection of orthologs of the molybdenum transporter AtMOT1 associated with variations in Mo content in poplar leaves. Eleven genomic intervals were found significantly associated with changes in Mo content, containing 44 candidate genes, of which 6 are predicted to have a transporter activity or can be related to ion homeostasis (Fig. 5a). On chromosome 6, twelve contiguous SNPs associated with variations in Mo were detected in the vicinity of PtrMOT1-1 (Potri.006G246000) and PtrMOT1-2 (Potri.006G245900), which share homology with a known molybdenum transporter AtMOT1 (Tomatsu et al., 2007; Tejada-Jiménez et al., 2007; Baxter et al., 2008b; Tejada-Jiménez et al., 2013; Fig. 5d). Nine SNPs were detected in the promoter region (295 to 2884 bp upstream the ATG start codon) and 3 SNPs were detected in the coding sequence (2 in the single exon and one in the 3’ UTR) of PtrMOT1-1 (Fig. 5b). At position Chr06:25345821, the missense mutation was associated with significant differences in accumulation in Mo, with the variants carrying the G allele and having a significantly lower Mo content than the variants carrying the T allele (Fig. 5c). PtrMOT1-1 harbored the highest transcript level across the 8 poplar tissues, in comparison to its paralog PtrMOT1-2, with a preferential expression in the roots, suggesting that PtrMOT1-1 is the functional copy of MOT1 in *P. trichocarpa*. Interestingly, the mutation at position Chr06:25345821 (highlighted in red in the alignment in Fig. 5d) caused the substitution of S (reference sequence) by an A in the protein sequence of PtrMOT1-1, which is identical to the sequence of its paralog MOT1-2, which does not appear to be the dominant copy of MOT1 in poplar. When considering the top 500 closest neighbors of PtrMOT1-1 in the co-expression network, this copy of MOT1 was found co-expressed with genes significantly enriched in genes involved in primary metabolism, which is consistent with previous reports on the function of AtMOT1 as cofactor for enzymes involved in several essential biological processes (Peng et al., 2018; Fig. 5f).

**Figure 5.**
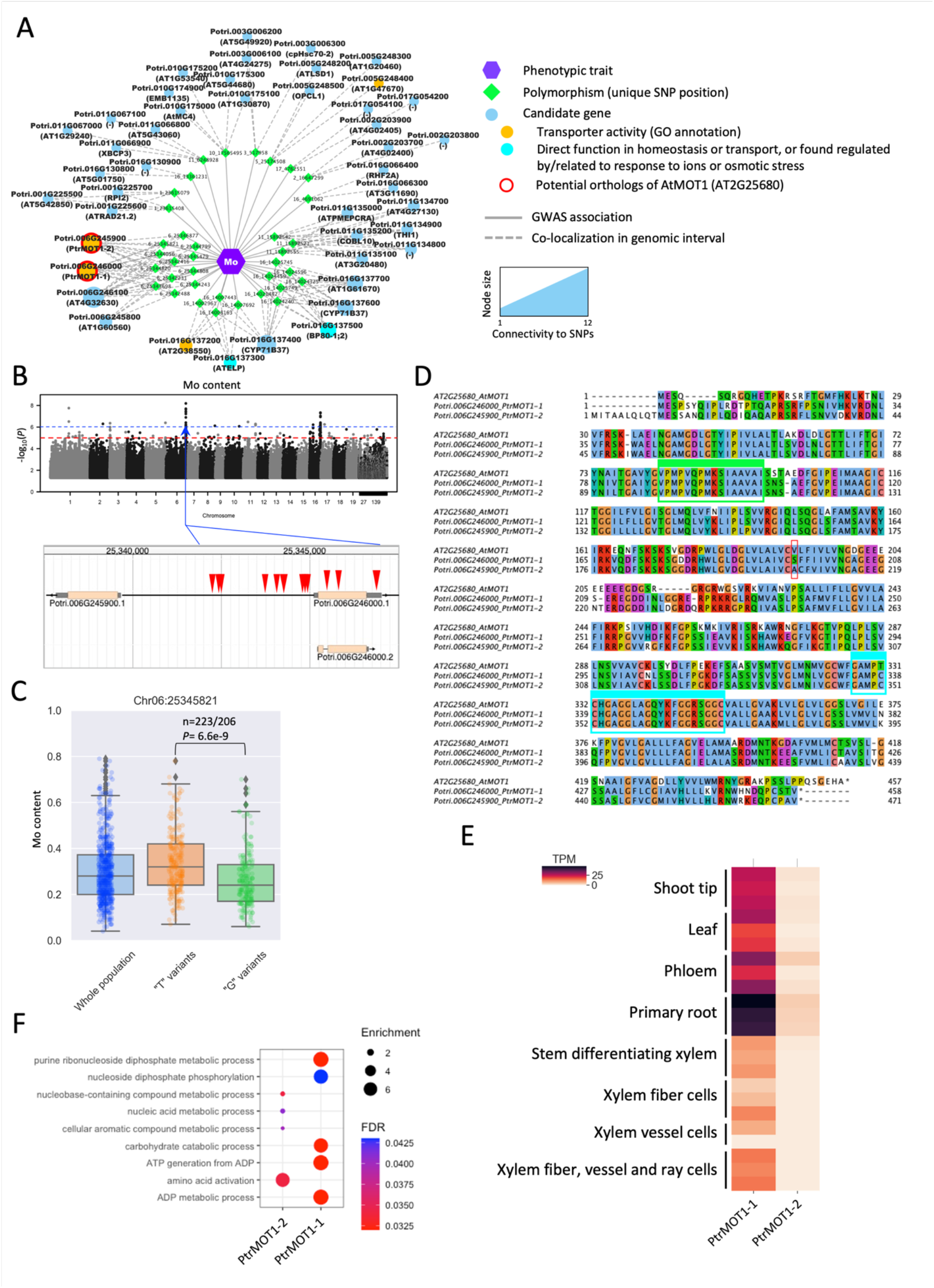
Detection of MOT1 as a determinant of Mo content in the leaves of poplar. **A**) Results of the GWAS for Mo content – all candidates identified by significant SNPs at a threshold of -log10(*P* value)>6 are represented and the two potential orthologs of AtMOT1 (Potri.006G246000 – PtrMOT1-1 and Potri.006G245900 – PtrMOT1-2) are highlighted in red (nodes represent either Mo content, SNP positions or potential candidate genes identified in the vicinity of the SNPs; edges represent a GWAS association or colocalization of the SNP and the annotated gene in the genomic interval). **B**) Manhattan plot for the Mo content (blue dotted line indicates the threshold for considering SNPs significant) and the genomic positions of the SNPs found significantly associated with variations in Mo content on chromosome 6. **C**) Difference in accumulation of Mo between the two alleles (T/G) – significance was assessed using a two-sided Student’s t-test. **D**) Protein alignment of *Arabidopsis* AtMOT1 (AT2G25680) and the two putative orthologs (PtrMOT1-1 and PtrMOT1-2), showing a conservation of the two domains previously found conserved in MOT1 across Eukaryotes and Prokaryotes (highlighted in green and blue). **E**) Transcript levels (RNA-seq) of PtrMOT1-1 and PtrMOT1-2 across different tissues of *Populus trichocarpa*. **F**) GO-BP enrichment results for the top 500 genes co-expressed with PtrMOT1-1 and PtrMOT1-2.

Next, to identify which biological processes underlie variations in elemental concentrations in poplar leaves, we relied on a co-expression analysis and the principle of ‘guilt-by-association’ to infer the biological function of some of the potential candidates. A co-expression approach was taken to identify the genes that are the most highly co-expressed with the candidate genes identified for each element and the main biological functions enriched in that group of genes was determined using a GO enrichment test.

Candidates were found to be co-expressed with genes enriched in numerous biological functions (Supplementary Data S4), with an over-representation in photosynthesis-related, transport-related, and cytoskeleton-related GO terms. Notably, 57 candidate genes showed a significant enrichment in multiple transport-related GO terms, including 10 that have transport activity or are involved in chemical homeostasis according to molecular function and biological process GO categories annotations (Fig. 6, Supplementary Data S4). For a total of 96 SNP positions, at least one of the flanking genes in the genomic interval was supported by significant co-expression with genes related to transport (Fig. 6; Supplementary Data S4). Examples include multiple genes related to P starvation response in *Arabidopsis* such as Potri.002G257800 (AtPHR1), Potri.008G130400 (AtAST91) and Potri.002G014900 (AtSEN1), that we found associated with micronutrient and trace-element content in poplar leaves. Potri.015G085100, the homolog of the nitrate transporter AtNRT3.1 was also found to be co-expressed with numerous genes enriched in functions such as ‘ion transport’, ‘cation transport’ and ‘transmembrane transport’ (Fig. 6).

**Figure 6.**
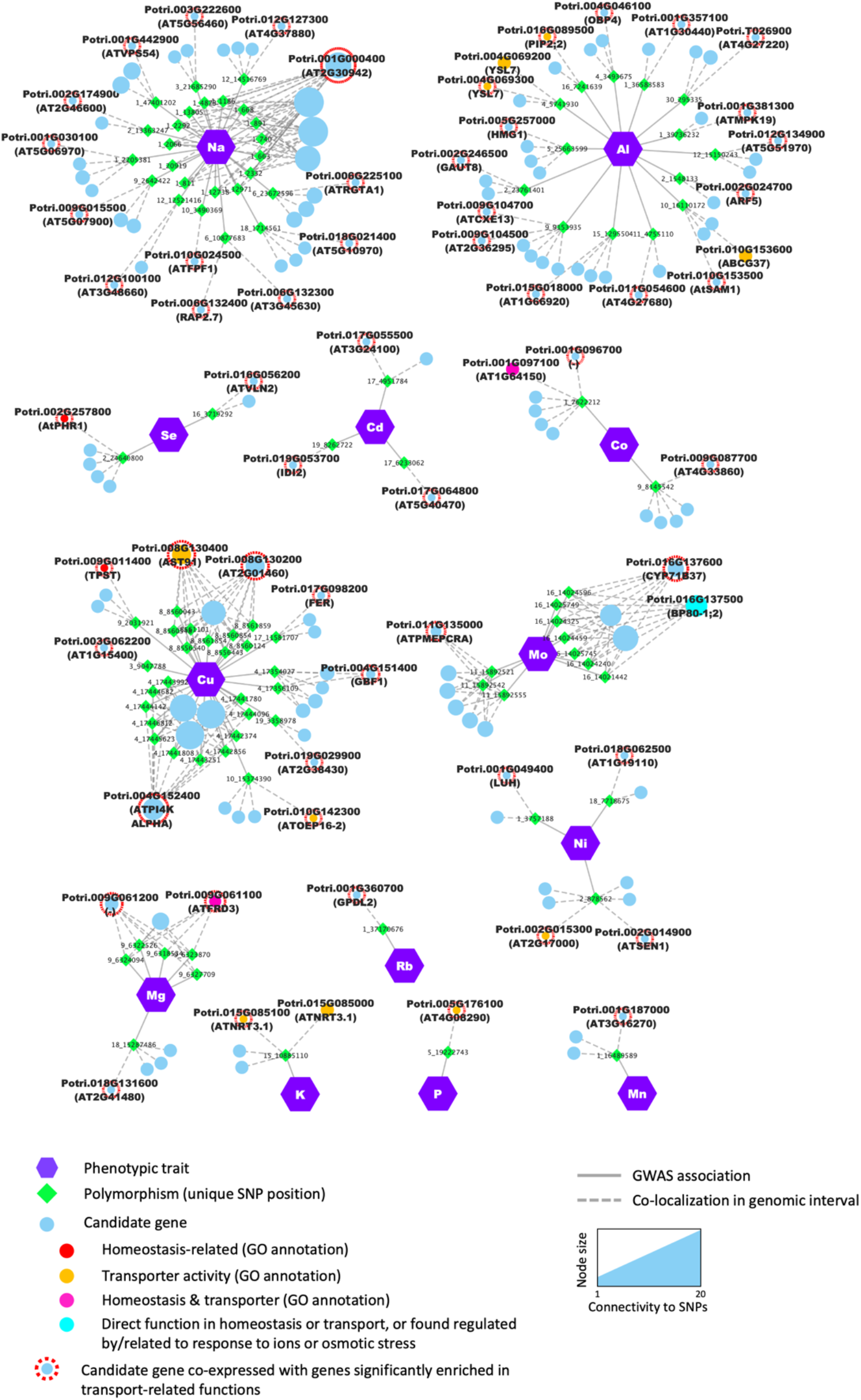
The GWAS identified candidate genes that are co-expressed with genes involved in ion homeostasis and transport. Nodes (i.e., colored circles and polygons) represent either phenotypic traits (i.e., mineral elements), polymorphisms (i.e., unique SNP positions) or potential candidate genes detected in the genomic interval flanking the significant SNP. Enrichment in genes related to transport among the co-expressed genes was tested using a GO enrichment analysis. Candidate genes which were co-expressed with genes enriched in transport-related GO terms are highlighted with a red outline. The exact number of co-expressed genes considered for GO enrichment for each candidate is provided in Supplementary Table 4. Edges (i.e., lines) represent significant associations detected by the GWAS or the colocalization of the SNP and the annotated gene in the genomic interval.

The two candidate genes Potri.004G069300 and Potri.010G182200, were picked from two distinct loci associated to variations in Al content in our conditions (Fig. 6; Supplementary Data S4). Potri.004G069300 was found to be co-expressed with genes that showed a significant enrichment in the GO term ‘transmembrane transport’, which is consistent with its predicted function given that it is a putative homolog of AtYSL7 that belongs to a family of metal-nicotianamine transporters that contributes to uptake and partitioning of metals in plant tissues (Curie et al., 2009; Waters et al., 2006). Interestingly, Potri.010G182200, a putative homolog of ATWI-12/SAG20, a senescence associated gene in *Arabidopsis* (Weaver et al., 1998), was found to be co-expressed with genes related to nicotianamine biosynthesis (Supplementary Data S4). Nicotianamine is an important molecule involved in the uptake, delivery, and circulation of metals in plants (Takahashi et al., 2003; Curie et al., 2009).

The SNP at position Chr09:6324094, was associated with variations in Mg and is located in the 3’ region of Potri.009G061100, a MATE efflux family protein found to be co-expressed with genes related to ‘ion transport’, ‘cation transport’, and biotic stress response (Supplementary Data S4). The *Arabidopsis* ortholog of this gene, Ferric Reductase Defective 3 / Manganese Accumulator 1 (FRD3/MAN1, AT3G08040), has been reported in numerous studies related to elemental accumulation in plant including iron (Delhaize, 1996; Rogers and Guerinot, 2002) and zinc (Pineau et al., 2012), with a role in biotic stress response (Scheepers et al., 2020). Together these results demonstrate that our GWAS was able to identify several genes known to be involved in the control of the ionome in plants, as well as a number of genes not reported to be involved in these processes.

Across all GWAS associations, six SNP positions were significantly associated with variations in more than one element. The most significant association for the macronutrient P was for the SNP located on Chromosome 18, position 14277823, which exhibited the lowest *p*-value for that phenotype (*p*=1.20E-11) and was also associated with two other elements: Co (*p*=1.96E-09) and Cu (*p*=3.58E-08; Fig. 7). This genomic interval harbors three candidate genes: Potri.018G118400, encoding a leucine-rich repeat family protein, Potri.018G118500, encoding a homolog of the *Arabidopsis* protein S-acyl transferase AtTIP1, and Potri.018G118600, predicted to encode a 2-oxoglutarate and Fe (II) dependent oxygenase superfamily protein. Interestingly, the *Arabidopsis* homolog of Potri.018G118600, AT1G49390, was reported to be systemically regulated under phosphorus starvation (Thibaud et al., 2010), suggesting that Potri.018G118600 could underlie the association observed in this genomic interval. Our GWAS results also suggest that Potri.018G118600 could play a role in the homeostasis of and/or in the response to Co and Cu in poplar.

**Figure 7.**
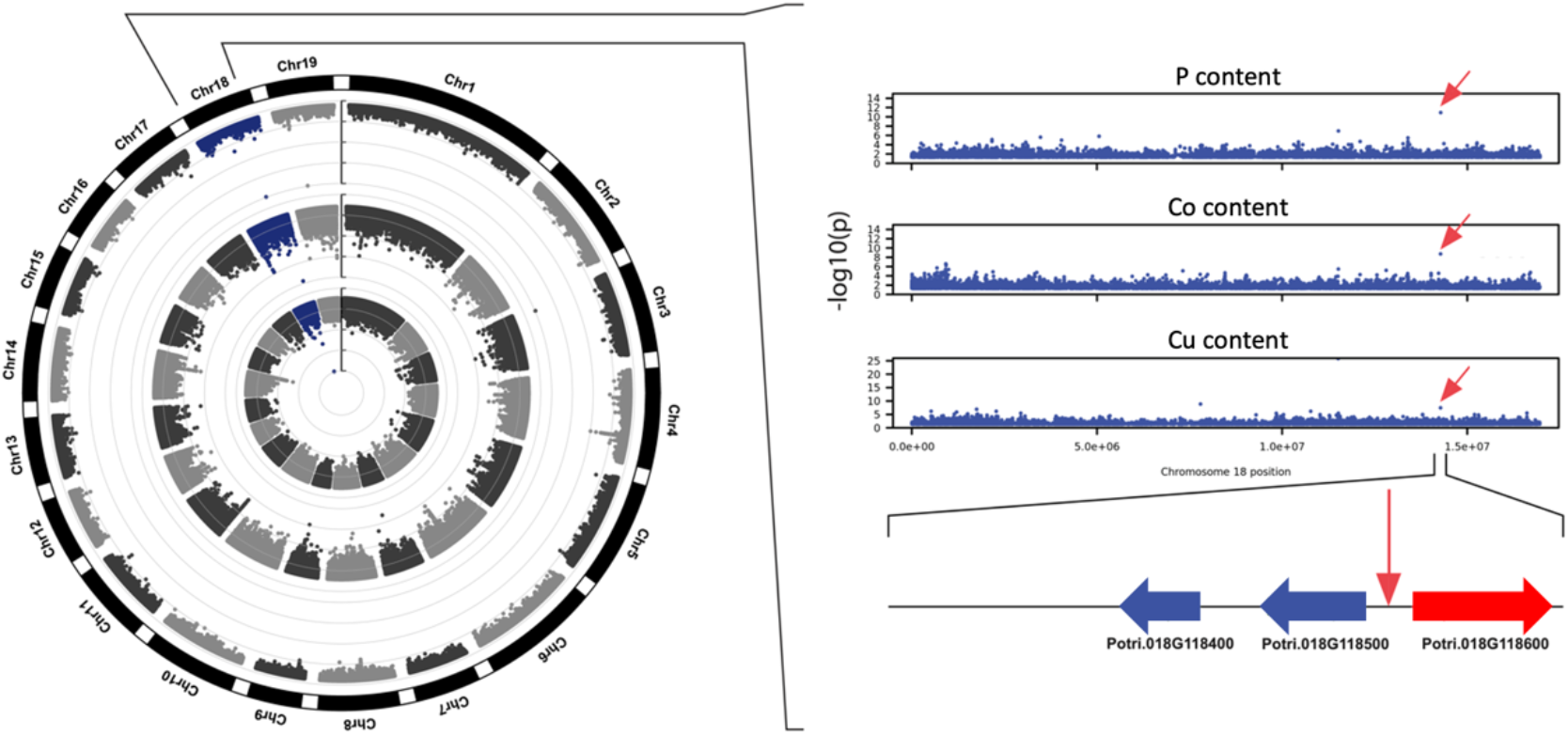
Circular Manhattan plot showing a genomic region on chromosome 18 exhibiting significant associations with abundance of elements P, Co, and Cu. Genome-wide associations detected for elements P, Co, and Cu are represented in the exterior, middle and interior rings of the circular Manhattan plot, respectively. Candidate gene Potri.018G118600 (red) was previously shown to be upregulated under phosphorus limitation.

Multiple SNP positions located in two genomic intervals were associated with Ca and Sr content (Fig. 8). On chromosome 4, the most promising candidate is Potri.004G184000, shares homology with *Arabidopsis* AT5G25940, which was identified by Schmidt et al. as a potential vacuolar membrane protein (Schmidt et al., 2007). P content was associated with 8 SNPs, all located in the same genomic interval on chromosome 14, of which 3 were also associated to S content. All SNPs pointed to one potential candidate, Potri.014G196600, predicted to encode a leucine-rich repeat protein kinase family protein with no clear function attributed yet (Fig. 8, Supplementary Data S4). These results suggest the existence of shared genetic regulation between elements that are essential nutrients for plant growth, and our approach uncovers several potential key players in this cross-homeostasis in poplar leaf tissues.

**Figure 8:**
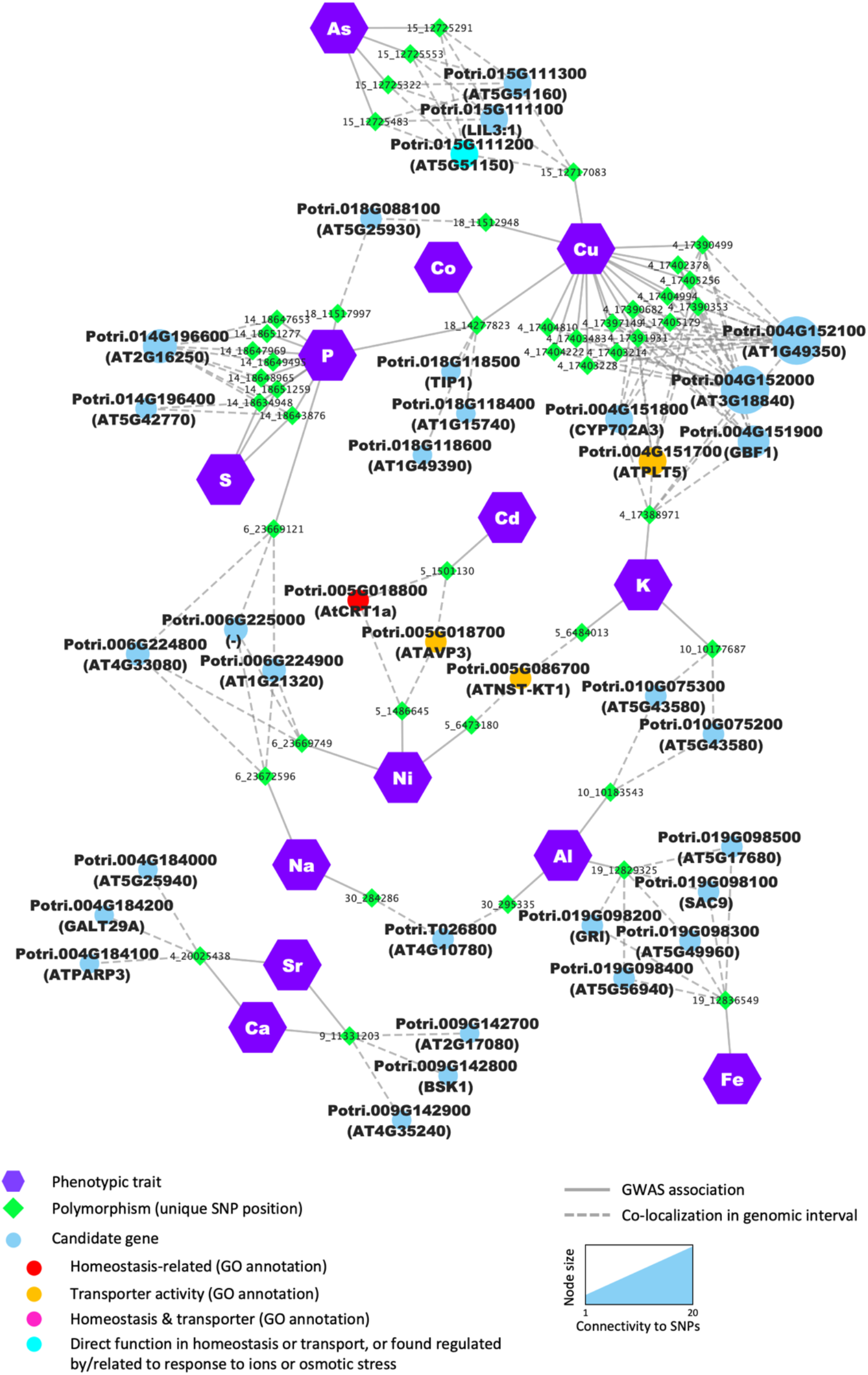
Several candidate genes are detected as determinants for multiple elements. Nodes (i.e., colored circles and polygons) represent either phenotypic traits (elements), polymorphisms (unique SNP position) or candidate genes detected in the genomic interval flanking the significant SNP. For candidate genes, node color indicates the different functional categories. Edges (i.e., lines) represent the significant association detected by the GWAS or the colocalization of the SNP and the annotated gene in the genomic interval.

In total, 34 genes were found associated with multiple mineral elements. These genes were identified based on SNPs directly associated with multiple mineral elements or located in the overlap between genomic intervals of different SNPs associated with multiple mineral elements, including 5 genes previously associated with ion homeostasis and/or with transporter activity, such as Potri.005G018800 and Potri.005G018700 both identified by two SNP positions found associated with the two heavy metals, Ni and Cd (Fig. 8, Supplementary Data S4). Potri.005G018800 is a homolog of AtCRT1a, a member of the calreticulin protein family, localized in the ER and involved in multiple cellular processes, including calcium homeostasis (Christensen et al., 2010). The second candidate, Potri.005G018700, is predicted to encode a H+-pyrophosphatase, homologous to *Arabidopsis* vacuolar pyrophosphatase AVP1. The function of AVP1 has been extensively characterized in multiple plant systems and has been shown to contribute to cation uptake and sequestration, with AVP1 overexpressing plants showing different levels of accumulation of K and Na in leaves and increased resistance to high NaCl or Mg and low P (Gaxiola et al., 2001; Yang et al., 2007, 2015). Interestingly, to date, none of these gene families have been directly related to the response to heavy metals.

In addition, Potri.004G151700, homologous to the broad substrate-specificity H^+^-symporter AtPMT5 in *Arabidopsis* (Klepek et al., 2010, 2005), was associated with both K and Cu content (Fig. 8; TableS4). Potri.004G151700 was found highly co-expressed with genes related to photosynthesis in our co-expression analysis, consistent with a critical role of K and Cu as macro- and micro-nutrients that impact photosynthetic processes. Several genes were identified as potential major regulators of ion partitioning for multiple elements. That was the case for Potri.018G088100, which was related to two elements critical for photosynthesis (P and Cu; Fig. 8) and was co-expressed with genes related to photosynthesis and primary metabolism (Supplementary Data S4). Other candidates were co-expressed with genes enriched in various biological processes, which highlights the tight connection between the elemental composition in the leaves and overall poplar physiology (Fig. 9; Supplementary Data S4).

In order to determine the extent genetic architecture controls of the ionome is conserved between a tree species and other plants, we tested the overlap between candidates found in our study and previous ionomics studies coupled with GWAS in rice and *Arabidopsis* (M. Yang et al., 2018). Despite differences in the experimental procedures and the limitations of orthology inference, 34 of the genes we detected as candidates in our study had potential orthologs associated to variations in elements in rice by Yang et al. (2018) (Supplementary Data S4). Notably, Potri.016G089500, predicted to encode a plasma membrane intrinsic protein subfamily PIP2, was associated with variation in Al and co-expressed with transport and ion homeostasis-related genes in poplar, and its potential ortholog (LOC_Os07g26660) was associated with variations in N content in rice (Supplementary Data S4; Yang et al., 2018). Furthermore, 7 overlapping candidates were also detected with the ionomics study that authors performed in *Arabidopsis*. Among these, the poplar gene Potri.T134000, a homolog of AtHK5, a high affinity K+ transporter in *Arabidopsis* (Rubio et al., 2008), was associated with Al content in our study and with Mo content in *Arabidopsis* (M. Yang et al., 2018). In addition to these results, the systematic detection of polymorphisms associated to Mo content located in the promoter and/or coding region of MOT1 across all three species (Supplementary Data S4), suggests a degree of conservation in the genetic architecture that controls variation in the ionome across plant species. Interestingly, most of the candidates overlapping across studies have not been related to ion homeostasis yet (Supplementary Data S4).

## 4. Discussion

During their life cycle, plants uptake mineral ions that are considered macronutrients, micronutrients, or trace elements. The uptake, transport, and storage of these ions within and across tissues are crucial processes that impact plant physiology. The paucity or excess amounts of some of these nutrients and trace elements can have detrimental effects on plant growth, leading to reduced yield, reduced fitness, and/or increased sensitivity to abiotic and biotic stressors (Pandey, 2015). Improving agronomic traits requires an understanding of how the ionomic profile varies and which genes mediate such variation. Several gene families are known to have conserved roles across plant species in these processes, including transporters, channels, and chelators (Aibara and Miwa, 2014); yet the genetic determinants of ionome variations remain largely unknown.

In numerous plant species, the use of natural genetic diversity has been used in forward genetics screening for identifying genes and regulatory mechanisms that underlie biological functions and determine a phenotype. By directly pointing at the genomic loci and the causal mutations, GWAS have been instrumental at identifying genes that are responsible for variations in many traits of interests in poplar (Muchero et al., 2015; Tuskan et al., 2018b; Xie et al., 2018; Zhang et al., 2018; Chhetri et al., 2020; Bryant et al., 2023; Nagle et al., 2023; Saint-Vincent et al., 2023). One major challenge for setting up this type of approach is the need to phenotype large populations (typically in the order of hundreds of individuals), with a method that can quantify a broad spectrum of elements with high sensitivity to capture slight variations across individuals and high accuracy.

Here we investigated three different platforms (LIBS, NAA, and ICP-MS) for the profiling of mineral elements in poplar leaf samples of a large natural population. All three methods agreed for several elements, however results for the quantification of K showed low correlation. Other elements such as Cu, Al, Na and Zn also showed relatively poor correlations between ICP-MS and NAA. This might be largely due to major differences in sample preparation, but it is also intrinsically linked to the analytical methods themselves. For NAA and LIBS analyses, pellets were pressed after humidification, while this pre-processing was not required for ICP-MS. The LIBS and NAA samples were pretreated by humidification which led to slightly different sample density which is manifested as weighing differences. Regarding the difference in the analytical methods, LIBS and NAA detect mineral elements on particles or small individual fragments of tissue (from pellets), which make them very sensitive to heterogeneity in the sample composition. In the contrary, IPC-MS works on large amounts of extracted material, which tends to make that method less sensitive to tissue heterogeneity. The low correlations among the three methods for K, compared to the other elements, suggests that K content may not have been uniform across the tested samples. The higher abundance in K in certain cell types, such as in the guard cells of the stomata for example, might contribute to variations in content measured through LIBS and NAA. Alternatively, ICP-MS values for K should be more accurate since a larger sample mass was dissolved for analysis than that taken for LIBS and NAA. Overall, NAA, LIBS, and ICP-MS are independent methods of analysis that each exploits different properties of the elements. NAA works on the nucleus and not the valence electrons, LIBS works on the energy level structure of the electron shells, and ICP-MS analyzes elements by ionization and mass separation. NAA is not sensitive to the valence state or chemical compound of the trace elements being measured which makes it less matrix sensitive and independent of chemical measurement methods such as ICP-MS. However, unlike ICP-MS, the detection sensitivity of NAA varies for each element according to the probability of neutron interaction and gamma-ray emission, gamma-ray energy, and decay half-life. These differences between the methods could explain the poor agreement between ICP-MS and NAA for Na.

Given that ICP-MS is the most suited platform to profile a broad range of mineral elements in a relatively high-throughput manner and considering that it has been largely used for ionome profiling in multiple plant systems (Baxter et al., 2012, 2008; M. Yang et al., 2018; Wu et al., 2021), we selected that method to profile the leaf ionome of 584 genotypes of *P. trichocarpa*. The analysis of the ionome of the population revealed that the content in micronutrients (Fe, Zn, Cu, Mn, B, Ni, Mo) and other trace elements (Cd, As, Co) was more variable than that of the macronutrients we quantified (P, K, Ca, Mg). A similar trend was observed in other species (e.g., rice and Arabidopsis; Yang et al., 2018), supporting the hypothesis that across annual and perennial plants, the concentration of the essential nutrients is under tight genetic control to ensure optimal growth, while micronutrients and trace elements are more prone to fluctuation.

To identify the genetic determinants of mineral ion composition in poplar leaves, we then performed GWAS, testing associations between a set of high-density SNPs (∼19 SNP per kb on average) and the elemental profile of 19 elements quantified using ICP-MS. This high-resolution mapping pointed to 922 unique potential candidate genes that could mediate, or be related to, variations in ion composition in the leaf tissues, of which 69 had putative orthologous genes already annotated to ion homeostasis and/or transport in other species. Even though the ionome of poplar leaves is not expected to be controlled exclusively by genes annotated with these GO terms, the significant enrichment in genes related to these biological processes or molecular function was expected and demonstrates that the GWAS captured relevant associations.

Our study provides strong evidence that a number of these candidates are the causal genes for the variation in elements. One ortholog of the high affinity molybdenum transporter MOT1 (Peng et al., 2018) emerges as the strongest candidate in our approach. In previous GWAS performed in *Arabidopsis* (Baxter et al., 2010; Chao et al., 2014, 2012; Forsberg et al., 2015), rice (M. Yang et al., 2018), maize (Wu et al., 2021) and cassava (Jin et al., 2022), MOT1 has been identified as a gene directly related to the Mo content. In our GWAS, one of the copies of MOT1 (PtrMOT1-1) showed a high degree of polymorphism, with several of these polymorphisms located in the promoter and coding region related to variations in Mo content. The fact that MOT1 is systematically identified as a strong candidate for Mo content in many ionomic GWAS suggests that Mo accumulation in plant tissues is highly dependent on MOT1 activity and reflects the level of conservation of that mechanism across species (Tomatsu et al., 2007; Peng et al., 2018), including in a tree species like black cottonwood poplar. The large number of SNP in the 5’ promoter region also suggests that the regulation of MOT1 is high controlled at the gene transcription level and that multiple transcription factors may exist that control transcription.

Another example of a direct association is the detection of PtrCHX16 (Potri.016G127800; Chanroj et al., 2012), a homolog of AtCHX26, as a candidate for Na content. Cation/H+ exchangers family (CHX) proteins belong to the MONOVALENT CATION PROTON ANTIPORTER-2 (CPA2) family of transporters involved in cation transport in plants (Chanroj et al., 2012; Khan et al., 2018). Although the exact function of most members of that family remains unclear, CHX were previously related to K+, Na+ and other monovalent cation transport and homeostasis and were shown to play a role in various cellular and physiological processes, including pH regulation, membrane trafficking, osmoregulation for stomatal opening and pollen tube growth (Cellier et al., 2004; Hall et al., 2006; Padmanaban et al., 2007).

Interestingly, another member of the CPA2 transporter family, PtrCHX25 (Potri.010G189100; Chanroj et al., 2012), a homolog of AtCHX21/23, together with Potri.010G100900 homolog of the Na^+^/H^+^ exchanger AtNHX7/SOS1 that belongs to the CPA1 transporter family, were associated with variations in Al content in poplar leaves. AtCHX21/23 was related to K transport, osmoregulation, and pollen tube growth, while extensive evidence demonstrated the role of AtNHX7/SOS1 in salt stress tolerance and K homeostasis (reviewed in Chanroj et al., 2012). This is the first report of this gene family being directly related to Al content. Similarly, the gene PtrNRT3.1C (Potri.015G085100; Plett et al., 2010) belongs to the NRT3 family and contributes to high affinity NO_3-_ transport in *Arabidopsis* (Plett et al., 2010), while it was found associated to K content in our study. Whitt et al. (2020) compiled a list of genes experimentally shown to change uptake, accumulation, and distribution of mineral elements across different species. Our analytical and computational approach related 9 genes of that list to variations in several elements in poplar (Supplementary Table S4). Except for the two copies of MOT1 that we related to Mo content, the 7 remaining genes were found associated to different elements than those reported in previous functional studies. Previous large-scale studies performed on the ionome of *Arabidopsis*, rice and maize (Baxter et al., 2010; Wu et al., 2021; M. Yang et al., 2018), uncovered many similar patterns of associations between an element and a likely candidate gene that was shown to be involved in the transport or homeostasis of another mineral element. Outside of the divergence between monocots and dicots (or annual *Arabidopsis* vs. perennial *Populus* within dicots), where in some cases, these associations can be explained by lineage specific evolutions that led to subfunctionalization of the paralogs, this type of indirect shared association highlights the importance of mineral interactions in plants.

The existence of strong relationships between different nutrients has been shown to underlie agronomic traits and physiological processes in many plant species (Baxter et al., 2015; Bouain et al., 2019). Typically, changes in content of one element can induce changes in the accumulation of other elements through competition for transport and uptake or by impacting absorption of other elements indirectly by changing developmental processes such as root architecture for example (Kellermeier et al., 2014) or cellular and physiological processes such as pH control and osmotic adjustment (Singh et al., 2013). A notable example is AVP1 in *Arabidopsis*, a pyrophosphatase (H+-PPase) proton pump that controls the electrochemical gradient across the vacuolar membrane. Overexpression of AVP1 improved NaCl tolerance in *Arabidopsis* (Gaxiola et al., 2001) and other plants including wheat (Brini et al., 2005), tobacco (Duan et al., 2007) and poplar (Yang et al., 2015). In *Arabidopsis*, overexpression of AVP1 resulted in enhanced cation uptake with an accumulation of Na and K in the leaf tissues, suggesting that AVP1 promotes sequestration of cations into the vacuole (Gaxiola et al., 2001). Interestingly it also influenced P nutrition, with AVP1 overexpressing *Arabidopsis*, tomato, and rice plants outperforming wild-type plants when grown with limited P. Overexpression of one of the related genes (PtVP1.1) in poplar also led to higher growth and resistance to NaCl treatment, together with a decreased amount of Na and higher amounts of K in poplar leaves (Yang et al., 2015). *Arabidopsis* plants overexpressing AVP1 also showed higher magnesium tolerance (Yang et al., 2018) as one of the phenotypes.

Many other examples of cross-homeostasis between nutrients exist in the literature, especially between the macronutrients N/P/K and metals or trace elements, such as Fe, Zn, S, Cu, Co, Mg and Rb (Baxter et al., 2008a; Bouain et al., 2019; Kocourková et al., 2020; Oliferuk et al., 2020). The significant correlation between profiles of multiple elements detected in our study was observed in other species (Neugebauer et al., 2018), which further supports the existence of interactions between mechanisms related to elements uptake and movement. In *Arabidopsis*, the multidrug and toxin efflux (MATE) transporter FRD3 is involved in distributing mineral elements across the plant tissues and has been shown to play an essential role in the cross-homeostasis of Fe and Zn. Mutants that have impaired growth typically accumulate high levels of different metals, such as Fe, Mn, and Zn in root and shoot tissues (Pineau et al., 2012; Roschzttardtz et al., 2011; Scheepers et al., 2020). It is therefore not completely surprising to find one poplar homolog of AtFRD3 associated with Mg content in our study. All of these observations reinforce the hypothesis that in attempts to identify regulators of the ionome, combinations of elements are likely to be the actual phenotype to consider for GWAS, rather than individual mineral elements, as proposed by Baxter (2015).

While previous studies highlighted the importance of considering interactions at the physiological, chemical, and genetic levels when studying mineral element homeostasis in plants, these interactions have remained largely unexplored due to their complexity. Our approach provides additional evidence of common genetic, and possibly physiological, mechanisms associated with mineral concentrations in poplar. Even though the potential causal gene for variations in one element was not systematically directly functionally associated with the homeostasis of that element, often, the likely causal gene was found co-expressed with genes involved in specific biological processes, such as photosynthesis, primary metabolism, and other essential biological functions, indicating that some of these elements interact at the physiological level. The identification of genomic loci associated to pairs of elements (i.e., K and Cu, P and Na, or the heavy metals Ni and Cd) illustrates direct genetic interactions between mineral element concentration, and as such, points to multiple new candidate genes potentially involved in the homeostasis of multiple mineral elements in plants.

## 5. Conclusions

By combining elemental profiling with high-resolution GWAS mapping across a population of *P. trichocarpa* genotypes, our study identified genomic regions and their associated mutations which putatively mediate variation in the ionome of poplar. Several of the genomic regions encompassed genes that have been previously reported to be involved in elemental accumulation in plants. Our approach identified strong candidate genes involved in the control of mineral ion content in poplar leaf tissues, such as the molybdenum transporter MOT1 that shows a high degree of conservation across plant species and is directly related to the Mo content. The results also revealed the existence of mineral nutrient interactions at the physiological and genetic level and provided candidate genes potentially involved in the cross-homeostasis of multiple elements in poplar. The detection of numerous key genes/proteins associated with ion uptake and circulation confirms that the leaf ionome profile is reflective of overall plant nutrient status. The combination of GWAS and ionomic profiling provides potential new target for engineering traits of interests, including enhanced elemental accumulation and salt stress resistance to promote poplar health and growth on marginal lands.

## Acknowledgement

This research was supported by the BioEnergy Science Center (BESC) and the Center for Bioenergy Innovation (CBI). BESC and CBI are Bioenergy Research Centers supported by the Office of Biological and Environmental Research in the US Department of Energy Office of Science. This research used resources of the Compute and Data Environment for Science (CADES) at the Oak Ridge National Laboratory, which is managed by UT-Battelle, LLC for the Office of Science of the U.S. Department of Energy under Contract Number DE-AC05-00OR22725.

## Supporting Information

**Supplementary Data 1**. Variation of the 20 elements profiled across the population of 584 *Populus trichocarpa* natural variants.

**Supplementary Data 2**. Results of GWAS at a threshold of -log10 (*p*) >6.

**Supplementary Data 3**. Number of significant SNPs detected through GWAS at a threshold of -log10(*p*)>6 for each element.

**Supplementary Data 4**. List of potential candidate genes flanking the SNPs identified by GWAS.

## Supplementary figure legends

**Supplementary Figure 1.**
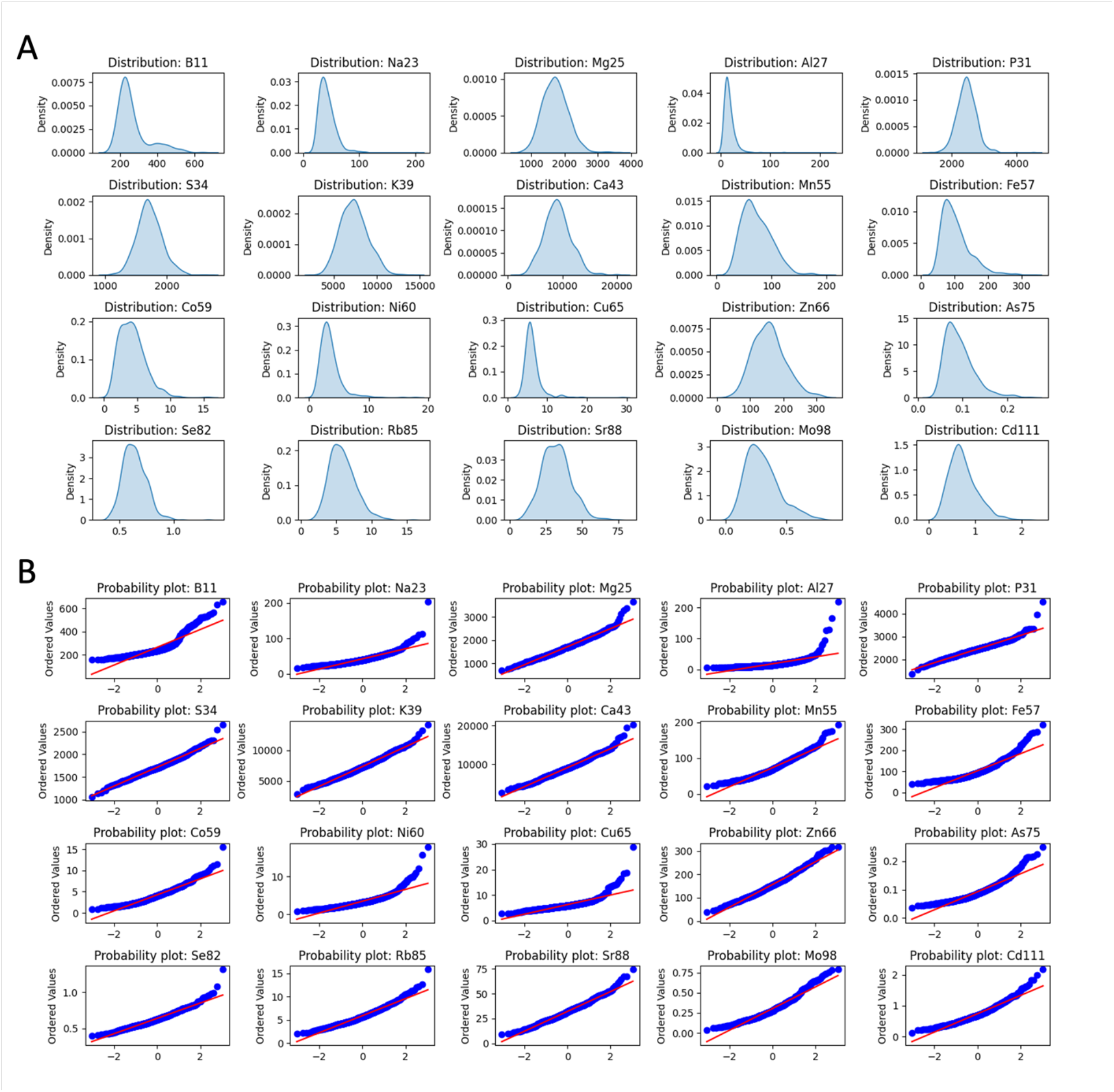
Distribution of the 20 elements across the leaf samples of 584 *Populus trichocarpa* natural variants. A) density plots and B) probability plots of the profiles.

**Supplementary Figure 2.**
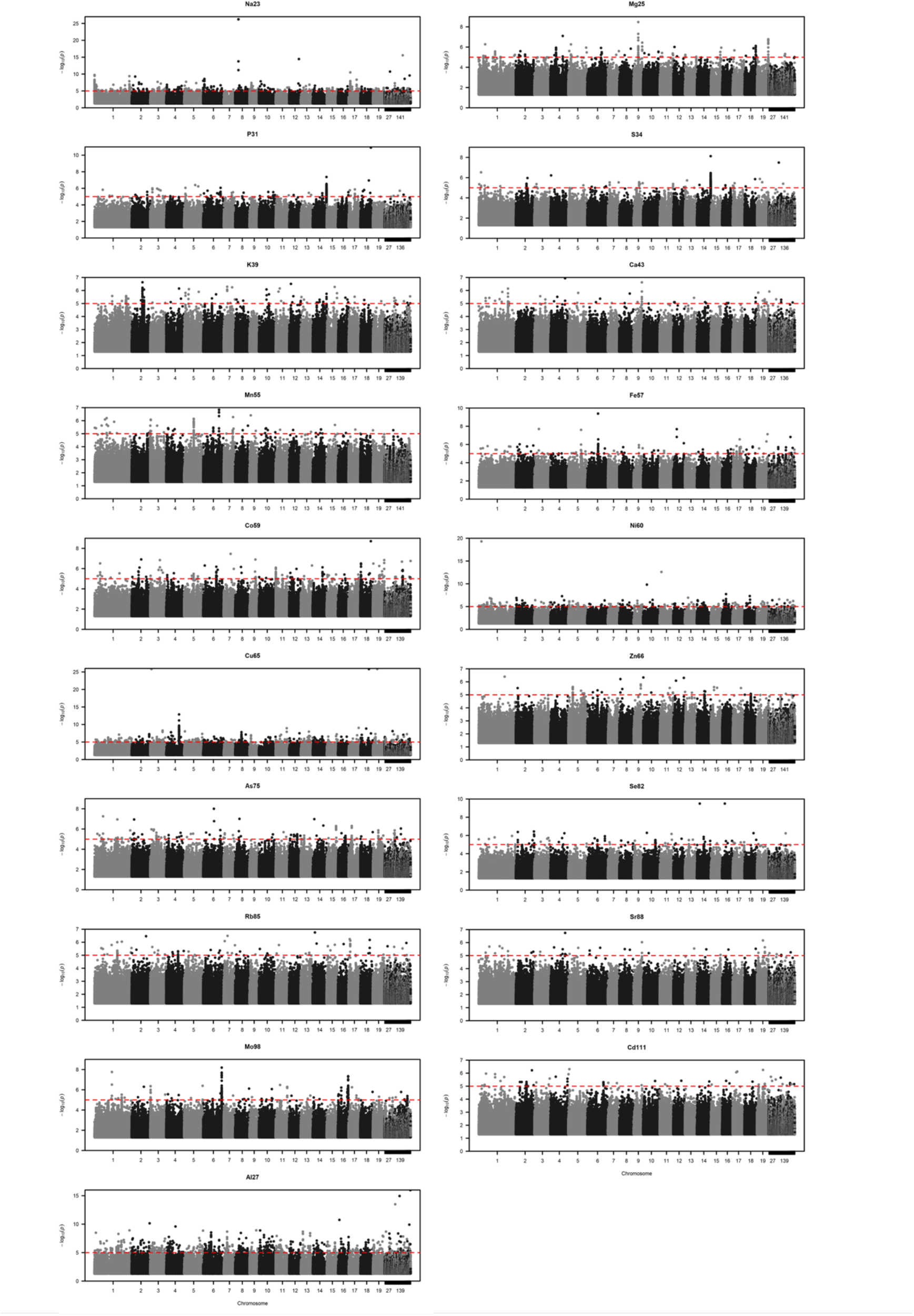
Manhattan plots resulting from the GWAS on the profile of 19 elements. B was not considered for the GWAS.

**Supplementary Figure 3.**
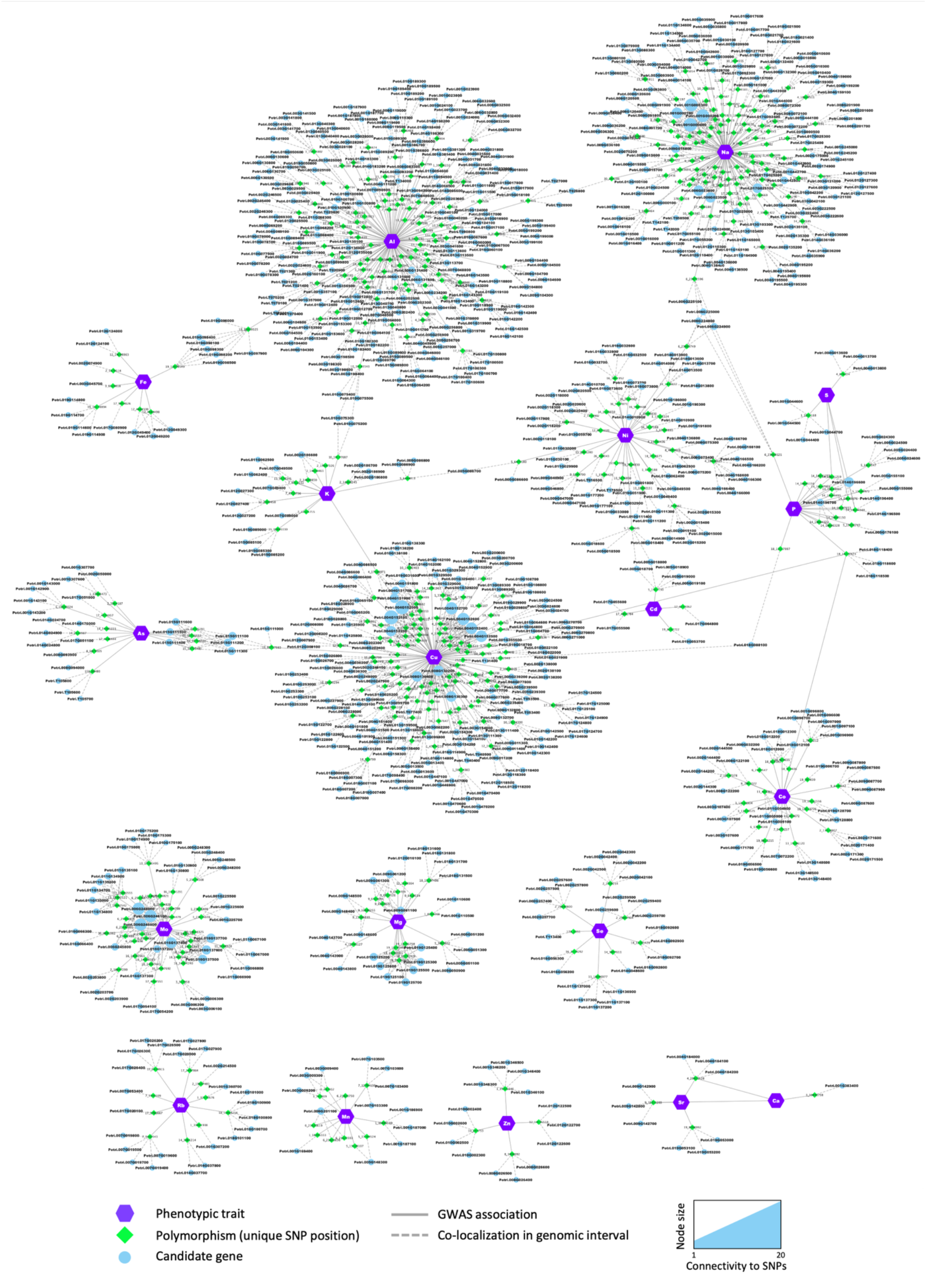
Network representation of the results of the GWAS. Nodes (i.e., circles and polygons) represent either phenotypic traits (mineral elements), polymorphisms (unique SNP positions) or potential candidate genes detected in the genomic interval flanking the significant SNP. For each candidate gene, the gene name of the best BLAST hit is provided in brackets in addition to the poplar accession number. When no *Arabidopsis* name was available, the gene accession number in *Arabidopsis* was provided. Edges (i.e., lines) represent the significant association detected by the GWAS (i.e., solid lines) or the colocalization of the SNP and the annotated gene in the genomic interval (i.e., dashed lines). For candidate genes, node color annotates the different functional categories and molecular functions.

